# ROMO1 and Mitochondrial Complex II/SDH are Required for Spare Respiratory Capacity and Glucose Homeostasis in Mice

**DOI:** 10.1101/2021.04.28.441811

**Authors:** Lisa Wells, Qian Yang, Caterina Iorio, Mithusha Peragerasingam, Zurie Campbell, Andy Cheuk-Him Ng, Courtney Reeks, Siu-Pok Yee, Robert A. Screaton

## Abstract

**Aims/Hypothesis:** Reactive oxygen species modulator 1 (ROMO1) is a highly conserved inner mitochondrial membrane protein that senses reactive oxygen species and regulates mitochondrial dynamics. ROMO1 is required for mitochondrial fusion *in vitro*, and silencing ROMO1 increases sensitivity to cell death stimuli. The physiological role of ROMO1 remains unclear.

**Methods:** To determine the role of ROMO1 *in vivo*, we used gene targeting in mice to ablate *ROMO1* in the whole mouse and to conditionally knock out *ROMO1* in the pancreatic beta cell. Mitochondrial functional analyses were performed on isolated mouse and human islets lacking *ROMO1*.

**Results:** We show that ROMO1 is essential for embryonic development, as ROMO1-null mice die before embryonic day 8.5, earlier than GTPases OPA1 or MFN1/2 that catalyze mitochondrial inner and outer membrane fusion. Knockout of ROMO1 in adult pancreatic beta cells results in impaired glucose homeostasis in young male mice due to an insulin secretion defect. Isolated islets from male, but not female, mice showed impaired glucose-stimulated insulin secretion. While mitochondria from female mice were morphologically normal, mitochondria in *<u>R</u>omo1* <u>a</u>dult <u>b</u>eta cell <u>k</u>nock<u>o</u>ut (RABKO) cells from male mice were swollen and fragmented, with a reduction in mtDNA content. Knockout of ROMO1 did not affect basal respiration in males or females, but deletion of ROMO1 in both sexes in mice and isolated human islets reduced spare respiratory capacity (SRC), which involved the specific loss of respiratory activity at Complex II/SDH. Aging of female ROMO1 KO mice resulted in loss of spare respiratory capacity and glucose intolerance.

**Conclusions/Interpretation:** Our data demonstrate that ROMO1 is a key regulator of mitochondrial bioenergetics and SRC and is required for effective nutrient coupling to insulin secretion in the beta cell. These observations point to a critical role for spare respiratory capacity in the maintenance of euglycemia and to the potential for targeting ROMO1-complex II to promote glucose coupling in settings of insulin insufficiency.

**Research in Context:** What is already known about this subject?

- ROMO1 is required for mitochondrial fusion
- Glucose coupling to insulin secretion is accomplished in part via generation of NADH during the oxidation of glycolytic metabolites in the TCA cycle
- Spare respiratory capacity is lost in aging

What is the key question?

- What is the physiological role of ROMO1 in the whole animal and the pancreatic beta cell?

What are the new findings?

- ROMO1 is essential for mouse development
- ROMO1 is required to maintain spare respiratory capacity (SRC) and to promote insulin secretion in the beta cells of mice and humans
- Ablation of the *Romo1* gene in the pancreatic beta cell leads to glucose coupling defects and glucose intolerance in young males and aged females
- Aging highlights the importance of SRC in the beta cell for maintaining euglycemia

How might this impact clinical practice in the foreseeable future?

- Aging is a significant risk factor for T2D. Human males, but not females, experience a loss of insulin secretion with age; designing strategies that enhance ROMO1 and complex II activity to promote SRC may help to reverse these effects.

## Introduction

Mitochondria are the primary organelles of ATP production from nutrient stores, and their integrity is central to cell survival. In the presence of oxygen, mitochondria harness electron-reducing equivalents derived from the tricarboxylic acid (TCA) cycle and fatty acid oxidation to fuel the electron transport chain and establish a proton gradient across the inner mitochondrial membrane (IMM) that drives ATP synthesis [1]. Mitochondria also participate in the synthesis and breakdown of macromolecules, including glucose, fatty acids, nucleotides, and amino acids, and serve as buffers for calcium ions [2, 3]. Mitochondrial dysfunction is a characteristic of aging [4] and is thought to underlie numerous human diseases, including cardiovascular disease [5], atherosclerosis [6, 7], neurodegeneration [8], as well as cancer and diabetes [9].

Mitochondrial dysfunction is most effectively monitored via functional assessment of mitochondrial oxidative capacity, and as they are critical sites for the generation and balancing of redox equivalents, by their production of reactive oxygen species (ROS) [10]. Accumulation of damaging ROS resulting from premature deposition of electrons on molecular oxygen is thought to drive mitochondrial dysfunction. ROS targets include mtDNA, proteins and membrane lipids, ultimately leading to reduced respiratory capacity. The uncontrolled production of ROS can induce permeabilization of the outer mitochondrial membrane (OMM) and the release of factors (cytochrome c, SMAC/Diablo) that activate the mitochondrial apoptotic cascade [11].

Reactive Oxygen Species Modulator 1 (ROMO1) is a highly conserved IMM protein reported to drive ROS production in cancer cells [12–15]. Furthermore, ROMO1 has been shown to govern the selective import of mitochondrial protein targets, including the protease YME-like 1 ATPase (YME1L) [16], and demonstrates nonselective cation channel activity *in vitro* [17]. In a genome-wide RNAi screen to identify regulators of mitochondrial dynamics, we showed that ROMO1 is a redox protein required for Optic Atrophy type 1 (OPA1) processing and that silencing ROMO1 increases ROS, fragments the mitochondrial network, and increases sensitivity to apoptotic stimuli [18]. In addition, U2OS cells lacking ROMO1 demonstrated a complete loss of spare respiratory capacity (SRC), a key feature of mitochondrial dysfunction under stress and in aging [19–22]. Despite the nearly 20 years since its discovery, the physiological role of ROMO1 is still unknown. To clarify the role of ROMO1 and of SRC *in vivo*, we generated mice lacking the *Romo1* gene in the whole animal and in the pancreatic beta cell.

## Methods

### Mice

All animal procedures were approved by the Animal Care Committee in Sunnybrook Research Institute, and experiments were carried out in accordance with the accepted guidelines. Animals were fed irradiated 18% Protein Rodent Diet (Envigo Teklad Global #2918-12159M)

#### ad libitum

Mice were in the modified barrier on a ventilated rack with automatic watering system. Allentown Polycarbonated Mouse cages (size 7.4” X 11.7” X 5.0 H) with corncob bedding made from ¼ inch 100% corncob, a nestlet made from pulped virgin cotton fiber and a PVC tube. Cage companion numbers ranged from 1-5. Mice were bred one male with one female in the same conditions, except each breeding cage had a Shepherd Shack made from unprinted, uncoated white book publisher-grade paper in place of the PVC tube. Temperature range was 20-21 °C, humidity was 60%, and light cycle was 12 hours light/12 hours dark. In all animal experiments, no statistical method was used to predetermine sample size and the experiments were not randomized. The investigators were not blinded to allocation during experiments and outcome assessment. All experimental groups were healthy prior to experimentation and had received only tamoxifen to induce *Romo1* gene recombination.

### Knockout Mice

The targeting vector designed carrying loxP sites flanking *ROMO1* exon 3 (Figure 1A) was transfected into C57BL/6-129/SvEv ES cells by electroporation. Exon 3 encodes Ile45-Cys79, which is predicted to encode the 10-residue loop between the first and second TM domain [18] and includes the TAA stop codon. The resulting recombined product is predicted to generate an out-of-frame mRNA subject to nonsense-mediated decay. G418-resistant colonies were screened by PCR. Targeted ES cells were injected into C57BL/6 blastocysts to create chimeric mice. Chimeric mice were bred with *Rosa26-Flpe* mice to remove the *PGK-neo* cassette. To create the null ROMO1 allele*, Romo1*^loxP/loxP^ mice (C57BL6/129sv background) were mated to universal Cre (HPRT-Cre) mice to generate cohorts as follows: *Romo1*^loxP/loxP^:HPRT-Cre^+^ and *Romo1*^loxP/+^: HPRT-Cre^+^, as well as appropriate controls lacking Cre. Genomic DNA was isolated using E.Z.N.A. Tissue DNA kit (Omega BIO-TSK D3396-02) using tail or ear clippings at 3 weeks of age, and PCR genotyping was performed with GoTaq polymerase (Promega M712). To detect the deletion of *Romo1* gene, the following primer sets were used: LoxgtF–5’TTACATTATCTGGCACGTCG and FrtgtR-5’ GCCTTTTAGCAGATAGCAAC. For WT mice there will be no product. For *Romo1* KO allele, the product is 500 bp. To detect the floxed-*Romo1* allele, the following primers were used: LoxgtF– 5’TTACATTATCTGGCACGTCG and LoxgtR– 5’AGGCTAGTATCGAACTCAGG. The WT allele product = 299bp, floxed allele product= 394bp. To detect Cre, the following primers were used: Cre 26 – 5’ CCTGGAAAATGCTTCTGTCCG; Cre 36 – 5’ CAGGGTGTTATAAGCAATCCC. Internal control primers were Gabra12: 5’-CAATGGTAGGCTCACTCTGGGAGATG-3’) and Gabra70 5’-AACACACACTGGCAGGACTGGCTAG-3’, product = 300 bp.

**1.**
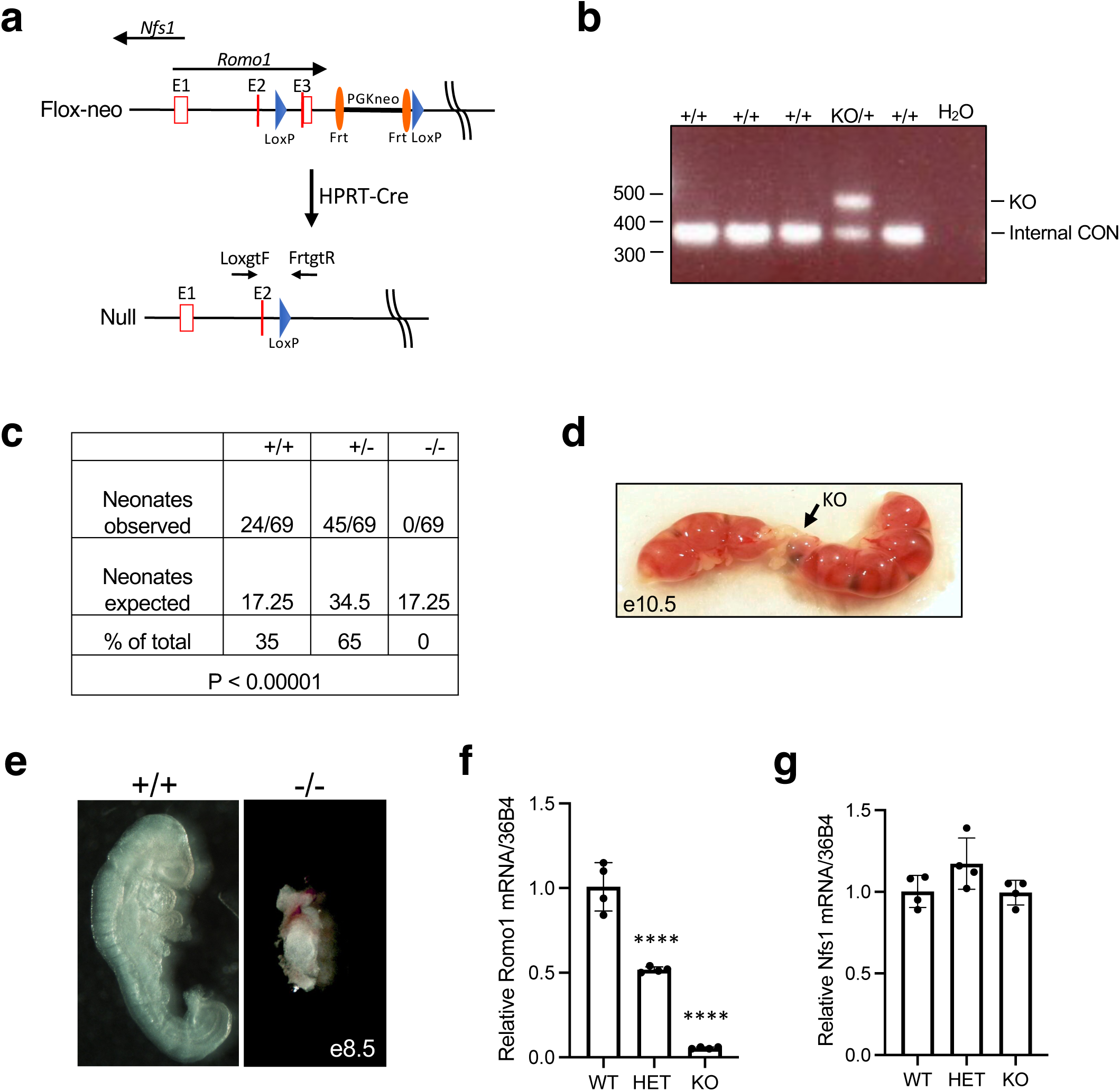
Early embryonic lethality in *Romo1* knockout mice. a. *Romo1* gene knockout strategy. Position of overlapping antisense gene *Nfs1* shown. Coding exons are indicated by solid red lines. Noncoding sequences are represented by red boxes. b. Genotyping of pups carrying KO allele. c. Live birth genotypes. d. Micrograph of embryos dissected at e10.5 then genotyped. KO embryo shown. e. Whole mount micrographs of WT and KO embryo at e8.5 showing early lethality of embryos lacking *Romo1*. f. RT-PCR analysis of *Romo1*, **** P-value: HET = 0.0004, KO = 0.00001. g. RT-PCR analysis of *Nfs1* levels from WT, HET, and cardiac *Romo1* KO (manuscript submitted), demonstrating *Nfs1* expression after Cre-mediated recombination is unperturbed. All data N.S.

### ROMO1 Adult Beta Cell Knockout (RABKO) Mice

Mice carrying the ROMO1 floxed allele were bred with mice carrying a *Pdx1*-Cre-ER^Tam^ transgene, which in adult mice drives tamoxifen-inducible Cre-mediated gene recombination only in the beta cell. 8-week-old mice were injected intraperitoneally (IP) with three injections of tamoxifen on alternating days at 5 mg/40g body weight [23].

### Histology and Immunohistochemistry

After timed breeding and confirmation of vaginal plugs, embryos were isolated at E8.5-10.5. For whole-mount immunohistochemistry, embryos were fixed in 4% paraformaldehyde in PBS overnight at 4°C prior to imaging by light microscopy. Paraffin-embedded pancreatic sections were stained with insulin (DAKO, A0564), and glucagon (DAKO, A0565) antibodies as described [23]. For β cell area determination, three sections per animal (n = 3 total animals per genotype) were stained with Insulin Ab (DAKO, A0564) and visualized with Vectastain ABC HRP Kit (Vector Labs, PK6100) and DAB solution kit (Vector Labs, SK4105). Slides were imaged at the Advanced Optical Microscopy Facility (AOMF), UHN Research. β cell area was determined using ImageScope software.

### Blood Glucose Measurements

Mice were fasted for 16 hours before sampling 30 uL of blood from the tail vein, then sampled again after 1 hour of refeeding with chow diet. Blood glucose levels were monitored using OneTouch Ultra Test Strips (AW07052001A).

### Glucose Tolerance Test

Intraperitoneal glucose tolerance testing (IPGTT) was performed as described [24]. Briefly, mice were fasted for 16 hours prior to IP injection of 2 g/kg (young mice) or 1 g/kg (aged mice) body weight glucose (SIGMA G7021) in PBS. Samples were taken at indicated times.

### Insulin Tolerance Test

Mice were fasted for 4 hours. Freshly prepared insulin (Humalog VL7510 DIN 02229704) was injected IP at 0.5 U/kg. 30 uL blood samples were taken at indicated times.

### Plasma Insulin Measurements

Mice were fasted for 16 hours. Blood samples from the lateral saphenous vein taken at indicated timepoints were transferred to heparinized capillary tubes (FisherBrand 22-260-950), clarified by centrifugation at 15,000g for 3 minutes and stored at - 80°C prior to assay with the Ultra-Sensitive Mouse Insulin Kit (Crystal Chem 90080).

### Arginine Tolerance Test

Mice were fasted for 16 hours. Freshly prepared L-Arginine (SIGMA, A8094) was injected IP at 1g/kg body weight, and 75 uL blood was collected in capillary tubes at 0, 2, and 5 minutes post-injection.

### Tissue Isolation

Mice were euthanized, weighed, and pancreas perfused with collagenase (Sigma C7657-5G) [23]. Islets were handpicked under microscope and cultured in RPMI media (Life Technologies, 11875-093) supplemented with 10% fetal calf serum (Wisent Bioproducts, 088150), 10 mM glucose and antibiotics at 37°C overnight before use. The left and right cortex and hypothalamus were collected and placed on ice, and stored at -80°C. 30 mg of each tissue was used for lysis with RIPA for western blot analysis.

### Glucose Stimulated Insulin Secretion (GSIS)

10 size-matched islets per sample were incubated in Krebs Ringer Buffer supplemented with 2.8 mM glucose for 2 hours, then treated with the indicated concentrations of glucose for 1 hour, followed by 45 mM KCl to evaluate depolarization-induced secretion [24]. Insulin content was determined by acid-EtOH extraction. Insulin levels were determined by HTRF using Insulin Ultra-Sensitive kit (Cisbio) [23]. The insulin levels were normalized to DNA levels using the Quant-iT dsDNA Assay Kit (Life Technologies).

### Cell Culture and Reagents

HEK293T and U2OS cells were obtained from ATCC and cultured in DMEM + 10% FCS + antibiotics in a humidified atmosphere with 5% CO_2_. HEK293T-17 cells were maintained below 90% confluence for no more than 20 passages. For plasmid transfection, HEK293T was transfected using PEI (Polysciences Inc). All cells were verified mycoplasma negative.

### Islet Culture

Islets were obtained from the NIDDK-funded Integrated Islet Distribution Program (IIDP islets) and the Alberta Diabetes Institute Research Islet Lab, Canada, lab of P. MacDonald (ADI IsletCore islets). Four male donors (aged 36 to 54 years old, BMI range 19.9 to 50.6 kg/m2) and four female donors (aged from 45 to 62 years old, BMI range 21.7 to 45.8kg/m2) without a prior diagnosis of diabetes were used to analyze mitochondrial function. Islets were cultured in non-tissue culture-treated petri dishes in PIM(S) media supplemented with 5% human AB serum, glutamine/glutathione and penicillin/streptomycin (all reagents from Prodo Laboratories Inc.).

### Antibodies

ERK2 (1:1000) and TOMM20 (1:1000) were from Santa Cruz, ROMO1 (1:2000) was from Origene, OPA1 (1:1000), SOD2 (1:2000) were from Abcam, and insulin and glucagon were from Dako. Insulin-647 conjugated antibody was from BD Biosciences. Antibodies were validated by RNAi or gene knockout in-house or based on literature.

**RNA interference (RNAi)** was performed as described [18].

### Lentivirus Production and Purification

Viral packaging (pCMV8.74, 20 ug/15 cm dish), envelope (pMD2G, 2.5 ug/15 cm dish) and pLKO.1 shROMO1 (20 ug/15 cm dish) were combined in 2 mL Optimem (Life Technologies) with PEI (Polysciences Inc) at a final concentration of 42.5ng/uL. The mixture was incubated for 15 min at room temperature and then added to 3×10^7^ HEK293T-17 cells in 20 mL DMEM containing 10% FBS and 50 IU/mL penicillin/50ug/mL streptomycin. Cells were incubated at 37°C 5% CO_2_ for 72 hr. Cell supernatant was then filtered through 0.2 uM filter and the fitrate centrifuged at 95,000xg for 2.5 hr at 4°C. Viral pellet was then resuspended in 200 ul PBS.

### SDS-PAGE and Western Blotting

Tissue samples were homogenized in RIPA buffer (50 mM TRIS pH 7.5, 150 mM NaCl, 0.1% SDS, 1% NP-40, 12 mM sodium deoxycholate) with Halt Protease Inhibitor Cocktail (Thermo Scientific) and 50 mM PMSF added fresh. Secondary antibodies (1:18000 IRDye 800CW Donkey anti-Rabbit IgG and 1:30000 IRDye 680RD Donkey anti-Mouse IgG for LI-COR; 1:3000 Goat anti-mouse/rabbit IgG (H+L) HRP conjugated antibodies (BioRad) for ECL detection) were diluted in 3% milk in PBS containing 0.01% SDS and 0.04% Triton-X 100. Immunoblots were scanned with Odyssey CLx Imaging System and quantified with LI-COR Image Studio Acquisition Software. For ECL detection, blots were treated with Amersham ECL Prime Western blotting detection reagent (GE Healthcare) and detected using HyBlot CL Autoradiography film (Denville Scientific). Western blotting to confirm ROMO1 KO was performed two weeks post-tamoxifen injection using 75 isolated islets per sample. Data are representative of at least two independent experiments.

### Quantitative PCR

DNA from islets isolated from male and female wild type and RABKO mice was extracted using DNeasy Blood & Tissue Kit (Qiagen). 20ug of DNA was used for each sample and subjected to qPCR using KAPA SYBR FAST qPCR Kit (Kapa Biosystems) on the Mastercycle Realplex system (Eppendorf). Relative levels of mitochondria genes Dloop1, Dloop2, Dloop3, CytB, 16S and Nd4 were analyzed and normalized to nuclear gene Tert using the 2^-ΔΔCt^ method using oligo sequences as described [25]. Data are from three biological replicates.

### Quantitative RT-PCR

Assays were performed on a Mastercycler EP Realplex 2 (Eppendorf) using a QuantiTect SYBR green PCR kit (Qiagen). Relative gene expression was determined by ΔΔCT normalized to expression of 36b4 internal control. The following primer sets were used: mouse *Romo1* (5’-GCCTTGTACCTCGACTCTGC-3’ and 5’-CCCCCAAGTCAGGTGTTCTA- 3’), mouse *Nfs1* (5’-GCAGCTCACAACCCCATTGTG-3’ and 5’- GGACTTCAGGACACCGCATC-3’), mouse *36b4* (5’-CCACGAAAATCTCCAGAGGCAC-3’ and 5’-ATGATCAGCCCGAAGGAGAAGG-3’).

### Mitochondrial Stress Test

Cells: U2OS cells were infected with lenti-shRNA targeting ROMO1 or non-silencing control on Seahorse XFe96 cell culture microplate on day 0 [18]. On day 3, cells were starved in 180 uL of Seahorse XF assay medium containing 1 mM glucose for 60 min, and oxygen consumption rate (OCR) was measured following treatment with stimuli as follows: 5 mM succinate, 5 mM glutamate, 2.5 mM malate, 5 mM pyruvate, 2 mM ADP, 2 uM oligomycin, 2 uM FCCP, 1 uM rotenone, 1 uM antimycin. For primary cells, islets isolated from mice or from human donors were washed in PBS and dissociated with Accutase (1mL/1000 IEQ) for 3-5 min at 37^°^C and triturated every 60 sec. Dissociated islet cells were seeded at a density of 60,000 cells/well in a XFe96 well Seahorse plate that was coated with poly(D)lysine. On the day of assay, cells were treated as described for U2OS cells and OCR was measured with stimuli as follows: 2 uM oligomycin, 1 uM FCCP, 1 uM rotenone, and 1 uM antimycin, and analyzed using an XFe96 Excellular Flux Analyzer (Agilent Technologies). For complex II assay, the dissociated cells were equilibrated with Mitochondrial Assay Solution (MAS) (Rogers G., etal. 2011) in CO2-free incubator for 1 hour, prior to injections with working concentrations as follows: Port A, rotenone (0.5 uM) and glucose (1 mM); Port B, succinate (10 mM final), saponin (25 mM), ADP (1 mM) and rotenone (0.5 uM); Port C, antimycin A (0.25 uM) and rotenone (0.5 uM). Protein content was quantified by BCA assay (Thermo Scientific). The results were normalized with Wave Edit Normalization Mode (Agilent Technologies). All assays were performed with biological triplicate and analyzed with Prism Version9.1.2 (GraphPAD).

### Electron Microscopy

Islets from control (CON) and RABKO mice were fixed with 4% paraformaldehyde, 1% glutaraldehyde for 24 hours, then washed with 0.1 M phosphate buffer, pH 7.0, and post-fixed with 1% osmium tetroxide in 0.1 M phosphate buffer, pH 7.0, for 1 hour. Islets were washed with 0.1 M phosphate buffer, pH 7.0 then dehydrated in an ethanol series. Islets were infiltrated with Embed 812/Araldite resin and cured at 65°C for 48 hr. Resin blocks were trimmed and polished and 90 nm thin sections were sectioned with a Leica Reichert Ultracut E ultramicrotome and mounted on TEM grids. Thin sections were stained with 5% uranyl acetate and 5% lead citrate. Sections were imaged using Transmission Electron Microscopy (Thermo Scientific Talos L120C) using a LaB6 filament at 120kV. For each sample, 20 grid squares were imaged at 3,400x and 13,500x magnification.

### Statistical Analyses

All studies were performed on at least three independent occasions. Results are reported as mean ± SEM. Statistical significance for animal experiments was determined as follows: for fasted and fed analyses, unpaired 1-way ANOVA was applied. For GTT and ATT assays 2-way ANOVA was used. All other data were evaluated using two-tailed unpaired Student’s *t*-test. All graphs were created using GraphPad Prism 6.0 (San Diego, CA) and analyzed statistically using SPSS 20 (IBM Corp., NY, NY), with significance accepted at p<0.05 (*), p<0.01 (**), p<0.001(***), p<0.0001(****).

## Results

### ROMO1 is essential for embryonic development in mice

To investigate the *in vivo* function of ROMO1, we generated complete and conditional mouse knockouts using a floxed ROMO1 allele (Figure 1a). As the *Nfs1* gene is transcribed from the opposite strand in a genomic region that overlaps with exon 1 of *Romo1*, we chose to target exon 3 with *loxP* sites. We crossed *Romo1^+/fl^* mice to mice carrying an *HPRT*-Cre transgene, consisting of Cre recombinase driven by promoter sequences from the HPRT gene that is involved in purine biosynthesis and is ubiquitously expressed. Following recombination, the predicted *Romo1* mRNA lacking exon 3 would generate an out of frame product and be subject to nonsense-mediated decay. To examine the effect of complete loss of *Romo1*, we confirmed the presence of the *Romo1* KO allele by PCR (Figure 1b) and analyzed the genotype of 69 progeny from crosses of heterozygous mice. *Romo1* heterozygous mice express a partially reduced level of ROMO1 protein and are viable (not shown). None of the progeny was homozygous for deletion of ROMO1, indicating that homozygous deletion of ROMO1 leads to embryonic lethality (Figure 1c). Dissection of embryos from pregnant female mice after timed breeding revealed that viable *Romo1^−/−^* embryos were not found after embryonic day 8.5 (E8.5; Figure 1d,e). We confirmed dose-dependent loss of *Romo1* mRNA in heterozygous and homozygous mice (Figure 1f). Expression of the *Nfs1* gene, which overlaps with *Romo1* but is transcribed from the opposite strand, was normal by RT-PCR (Figure 1g). Thus, we conclude that our knockout strategy specifically targeted *Romo1*, and that *Romo1* is essential for embryonic development in mice.

### ROMO1 is required for energy coupling in the pancreatic beta cell

Having established that ROMO1 is essential for embryonic development in mice, we next sought to test if ROMO1 is also required to regulate cellular energetics *in vivo*. Given that insulin secretion by a pancreatic beta cell is an energetically demanding process accompanied by significant ROS production that must be controlled by cellular reductive systems[26], we generated a pancreatic beta cell-specific knockout of ROMO1 using a *Pdx1*-*ERTam-Cre* transgenic line [23, 24], which after 8 weeks of age expresses Cre recombinase only in the pancreatic beta cell compartment of the islet (Figure 2a). By breeding *ROMO1^+/flox^* mice to *ROMO1^flox^*^/+^*:Pdx1-CreER^Tam^*mice, we generated *ROMO1^flox^*^/*flox*^*:Pdx1-CreER^Tam^*mice, that in the presence of tamoxifen, produce the nonsense ROMO1 transcript in adult beta cells that is then eliminated. We refer to these mice as <u>R</u>OMO1 <u>a</u>dult <u>b</u>eta cell <u>k</u>nock<u>o</u>ut (RABKO) mice. As controls, we used *ROMO1^+^*^/+^*:Pdx1-CreER^Tam^* mice (referred to below as CON). RABKO mice were born at normal frequencies and had body weights indistinguishable from littermate controls (not shown). Western blot revealed ∼70% reduction of ROMO1 protein in islets isolated from RABKO mice, consistent with the relative fraction of beta cells in islets, as well as unchanged levels of ROMO1 in the cortex and hypothalamus (Figure 2b), consistent with our previous work with the *Pdx1-CreER^Tam^* transgene [23, 24]. Islet morphology and insulin staining was normal in male and female RABKO mice (Figure 2c) with no change in islet size (Figure 2d). Immunostaining for insulin and glucagon confirmed that alpha cell populations were unaffected (Figure 2e).

**2.**
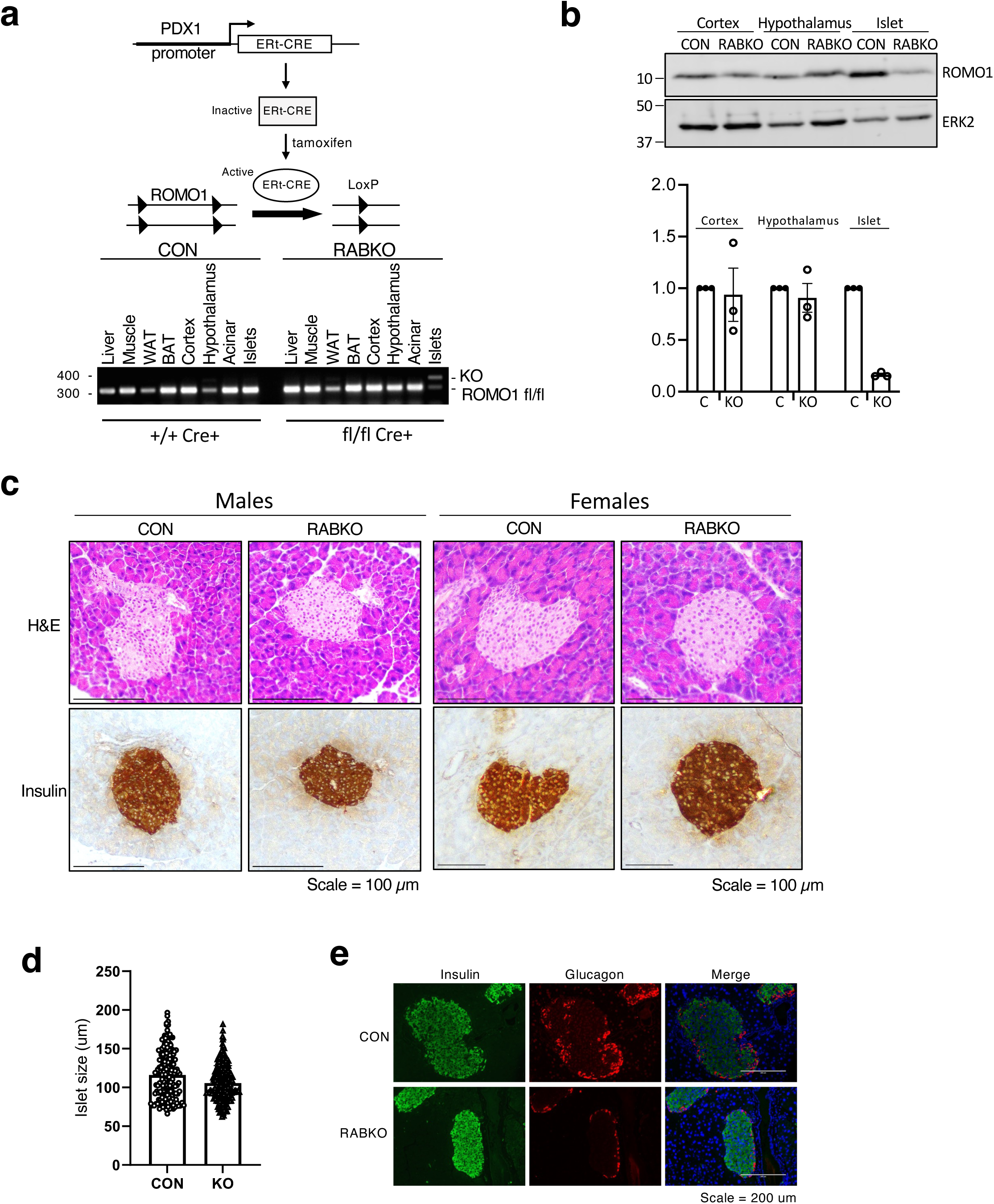
Generation of <u>R</u>OMO1 <u>A</u>dult <u>B</u>eta cell <u>K</u>nock<u>O</u>ut (RABKO) mice. a. Conditional *Romo1* adult beta cell knockout schematic and genotyping analysis of DNA from Cre+ animals carrying *Romo1* +/+ or fl/fl alleles, showing recombination only in fl/fl Cre+ islets and no other tissue. b. Western blot of ROMO1 in cortex, hypothalamus, and islets of CON and RABKO mice showing loss of ROMO1 protein only in the islet samples. Loading control for the samples is ERK2. c. (top) H+E staining of pancreatic sections from CON and RABKO mice. (bottom) Insulin staining in pancreatic sections. d. Quantification of islet size in male CON and RABKO mice. e. Immunofluorescent imaging of insulin (green) and glucagon (red) with DAPI counterstain on pancreatic sections from WT or RABKO mice showing normal islet morphology.

Male and female RABKO mice at 10 weeks of age were normoglycemic after an overnight fast but showed elevated blood glucose after 1 hour of refeeding, with RABKO males showing more pronounced hyperglycemia than females (Figure 3a,b). Heterozygous male and female mice were normal. Male but not female RABKO mice showed lower plasma insulin after 1 hour of refeeding (Figure 3c,d), and when challenged with an IP glucose bolus to evaluate acute insulin secretion capacity, male but not female mice were glucose intolerant (Figure 3e,f). The glucose intolerance in the males coincided with significantly reduced plasma insulin levels (Figure 3g,h). To evaluate beta cell responsiveness to a depolarizing stimulus *in vivo*, we next performed an arginine tolerance test. Male, but not female, RABKO mice failed to mount a significant acute insulin secretory response (Figure 3i,j). Taken together, these data indicate that ROMO1 in the beta cell is required for nutrient coupling and maintenance of euglycemia in male mice.

**Figure 3.**
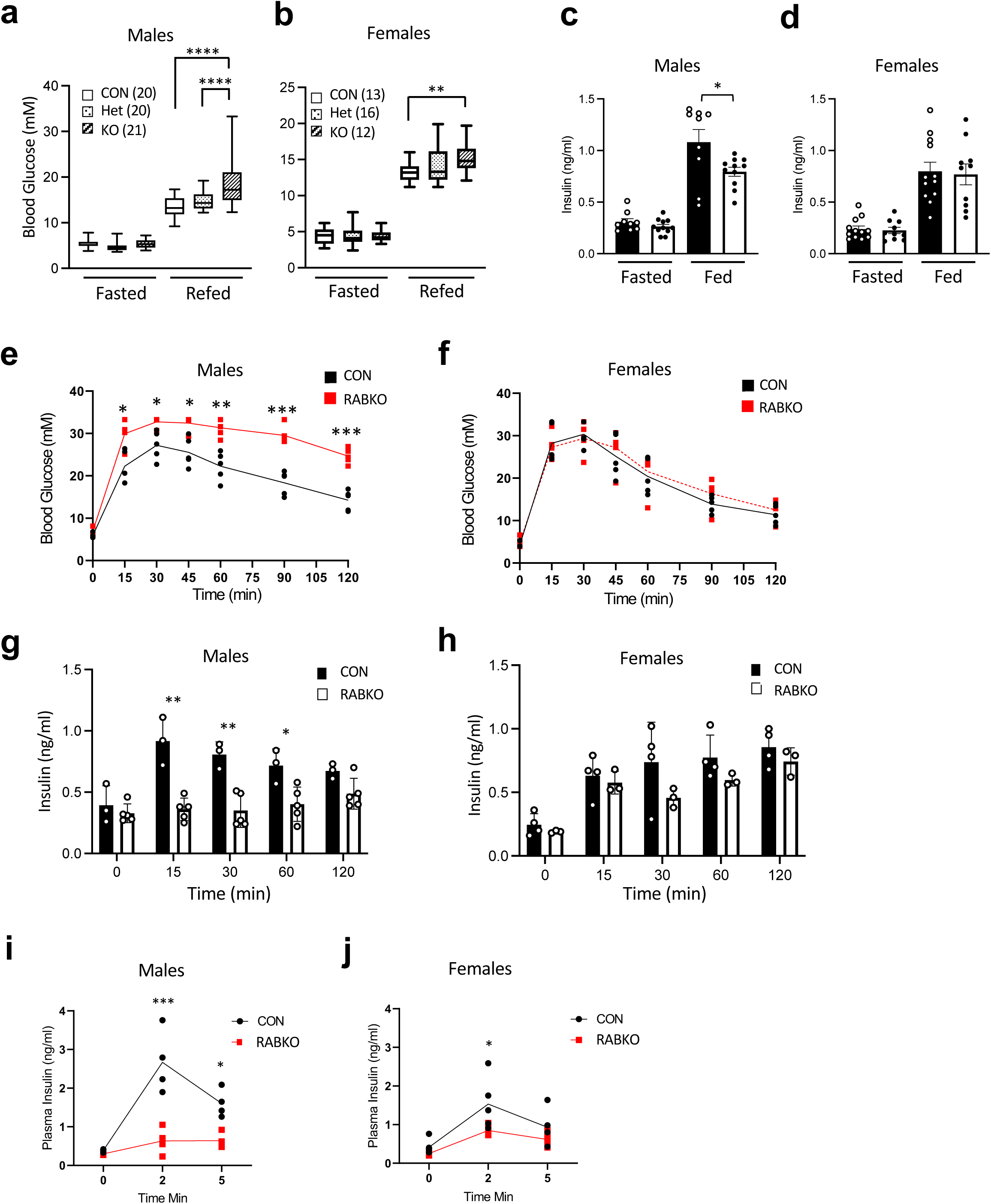
Glucose intolerance in RABKO mice. a. Fasted and 2-hour refed blood glucose in 10-week-old male *Romo1^+/+^*;*Cre^+^* (CON), *Romo1^+/fl^*;*Cre^+^* (Het) and *Romo1^fl/fl^*; *Cre^+^* (KO) mice. **** P-values = 0.0001. b. Fasted and 2-hour refed blood glucose in 10-week-old female *Romo1^+/+^*;*Cre^+^* (CON), *Romo1^+/fl^*;*Cre^+^*(Het) and *Romo1^fl/fl^*; *Cre^+^* (KO) mice. ** P-value = 0.0012. c. Plasma insulin levels from male mice shown in a. P-value = 0.029. d. Plasma insulin levels from male mice shown in b. e. IPGTT assay in CON and RABKO 10-week-old male mice following injection of 2g glucose per kg body mass. P-values = *15 min 0.025, *30 min 0.034, *45 min 0.025, **60 min 0.007, ***90 min 0.0008, ***120 min 0.0005. f. IPGTT assay in CON and RABKO 10-week-old female mice. All data N.S. g. Plasma insulin levels in CON (black bars) and RABKO (white bars) 10-week-old male mice during IPGTT glucose challenge from e. P-values = **15 min 0.0013, **30 min 0.0026, *60 min 0.02. h. Plasma insulin levels in CON (black bars) and RABKO (white bars) 10-week-old male mice during IPGTT glucose challenge from f. i. Plasma insulin levels in CON and RABKO male mice after arginine injection. P-values = ***2 min 0.0003, *5 min 0.014. j. Plasma insulin levels in CON and RABKO female mice after arginine injection. P-value = *2 min 0.03.

### Islet functional defects in male RABKO mice

Static insulin secretion assays using isolated islets from CON and RABKO male and female mice demonstrated that both glucose-stimulated and KCl/depolarization-induced insulin secretion was reduced by ∼ 50% in male RABKO islets compared to CON islets (Figure 4a), despite a normal insulin content (Figure 4b). We previously reported altered cristae in cultured cells silenced for ROMO1 by RNA interference [18]. To determine whether this effect also occurs in primary beta cells carrying a ROMO1 deletion, we analyzed mitochondrial ultrastructure in male RABKO beta cells. Mitochondria in beta cells from control mice appeared as long tubular structures with ordered cristae (Figure 4c, top panels). In contrast, mitochondria in RABKO beta cells displayed fewer cristae (Figure 4c, bottom panels, and Figure 4d), some of which lacked a clear connection to the IMM, as well as rod-like structures (not shown) seen in mitochondria defective in membrane fusion [27–29]. Furthermore, RABKO mitochondria were visibly swollen, and insulin granules in RABKO mice showed rod-shaped insulin crystals, as well as a reduction in electron density (Figure 4e), consistent with immature or pathological Zn^2+^-dependent packing of insulin [30]. Islets isolated from female RABKO islets were phenotypically normal, indicative of a sex-specific defect in glucose coupling and/or granule recruitment post-membrane depolarization (Supplementary Figure 2). Coincident with these alterations, islets from male but not female RABKO mice showed a ∼50% reduction in mtDNA content (Figure 4f).

**Figure 4.**
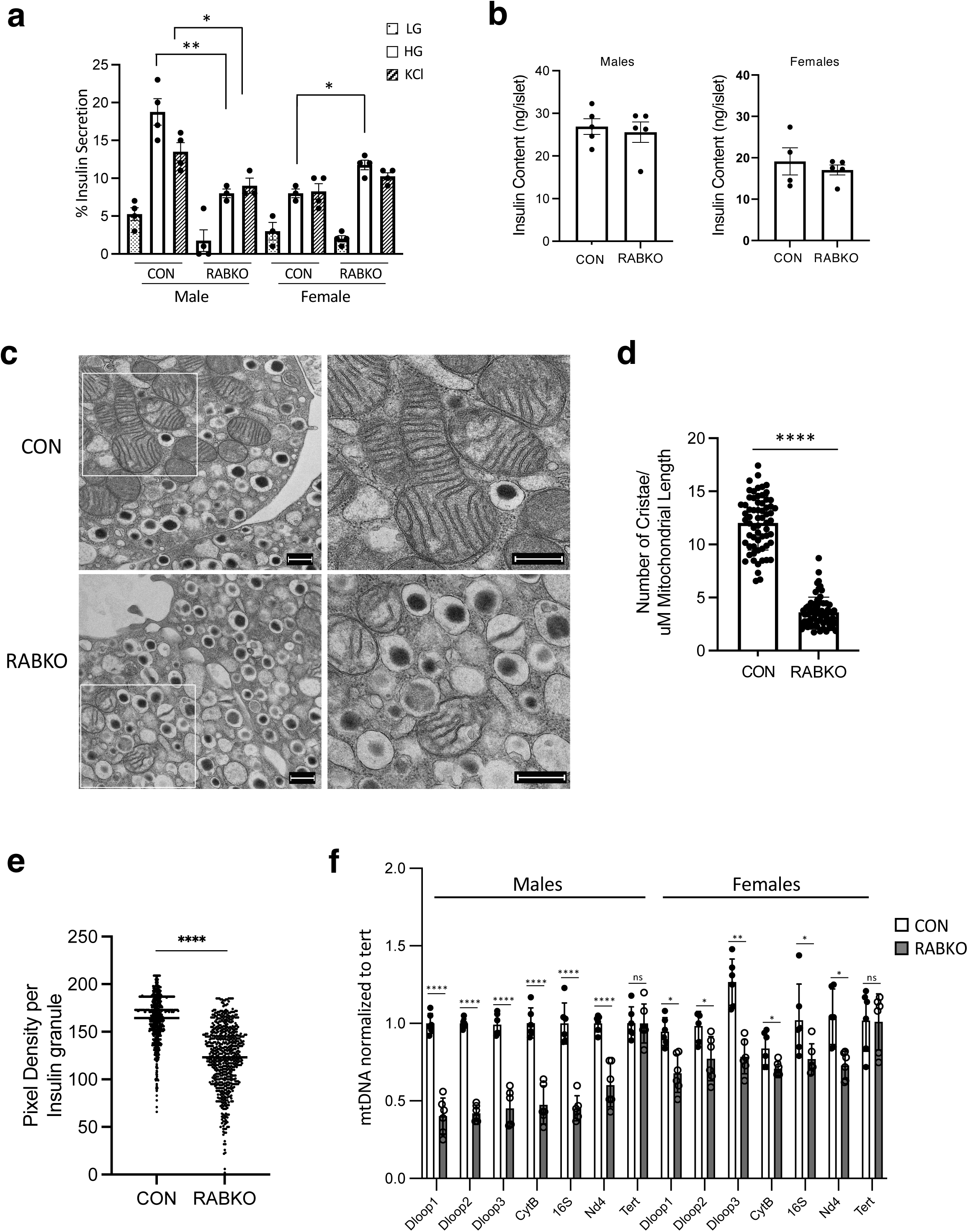
Ultrastructural alterations in mitochondria of male RABKO islets. a. Static glucose-stimulated insulin secretion (GSIS) assay with islets isolated from 10-week-old male and female CON and RABKO mice incubated with 2.8 mM glucose prior to stimulation with 16.7 mM glucose or 45 mM KCl to depolarize the cells. P-values: males **HG 0.0048, *KCl 0.029; females *HG 0.014. b. Insulin content in islets from CON and RABKO male and female mice. c. Electron micrographs showing mitochondrial morphology in 10-week-old RABKO mice. Images on right are 2-fold magnification of highlighted region (white box on left). Scale bars = 500 nM. d. Number of cristae per unit mitochondrial length in beta cells from male CON and RABKO mice. ****P-value = 0.0001. e. Violin plot of pixel intensity in insulin granules in male CON and RABKO beta cells. ****P-value = 0.0001. f. qPCR data of mitochondrial DNA in male and female CON and RABKO mice at 16 weeks of age. Relative levels of mtDNA genes Dloop1, Dloop2, Dloop3, CytB, mito16S, and Nd4 in islets from CON and RABKO male and female mice were normalized to level of the nuclear DNA gene, Tert. P-values as follows: * < 0.05, ** < 0.01, **** < 0.00001.

### Loss of Spare Respiratory Capacity in RABKO beta cells

To determine if these ultrastructural changes in male beta cells lacking ROMO1 resulted in loss of respiratory activity, we compared oxygen consumption rates in islets from male and female CON and RABKO mice. RABKO females, but not RABKO males, showed an increase in basal respiration rate (Figure 5a). However, the SRC, defined as the difference between maximal (post-FCCP treatment) and basal respiration rates, was reduced by ∼50% in both sexes in RABKO beta cells (Figure 5b), consistent with our previous work [18]. Proton leak was increased in islets from both females and males, consistent with a loss of efficient bioenergetic coupling in the absence of ROMO1 (Supplementary Figure 3). Interestingly, we noticed that the absolute values of SRC were consistently higher in islets from females, indicating female islets have a higher set point of SRC than males. After deletion of ROMO1, SRC in female RABKO mice was lowered to the value observed in control male mice. Silencing of ROMO1 in isolated human islets resulted in > 60% reduction in SRC in islets from male subjects (Figure 5c) and > 40% reduction in female subjects (Figure 5d), with total SRC in males being 30% lower than n females (Figure 5e). Taken together, these data indicate that ROMO1 is required to maintain SRC in male and female mice, and that islet cells in female mice and humans have an intrinsically higher SRC than males.

**Figure 5.**
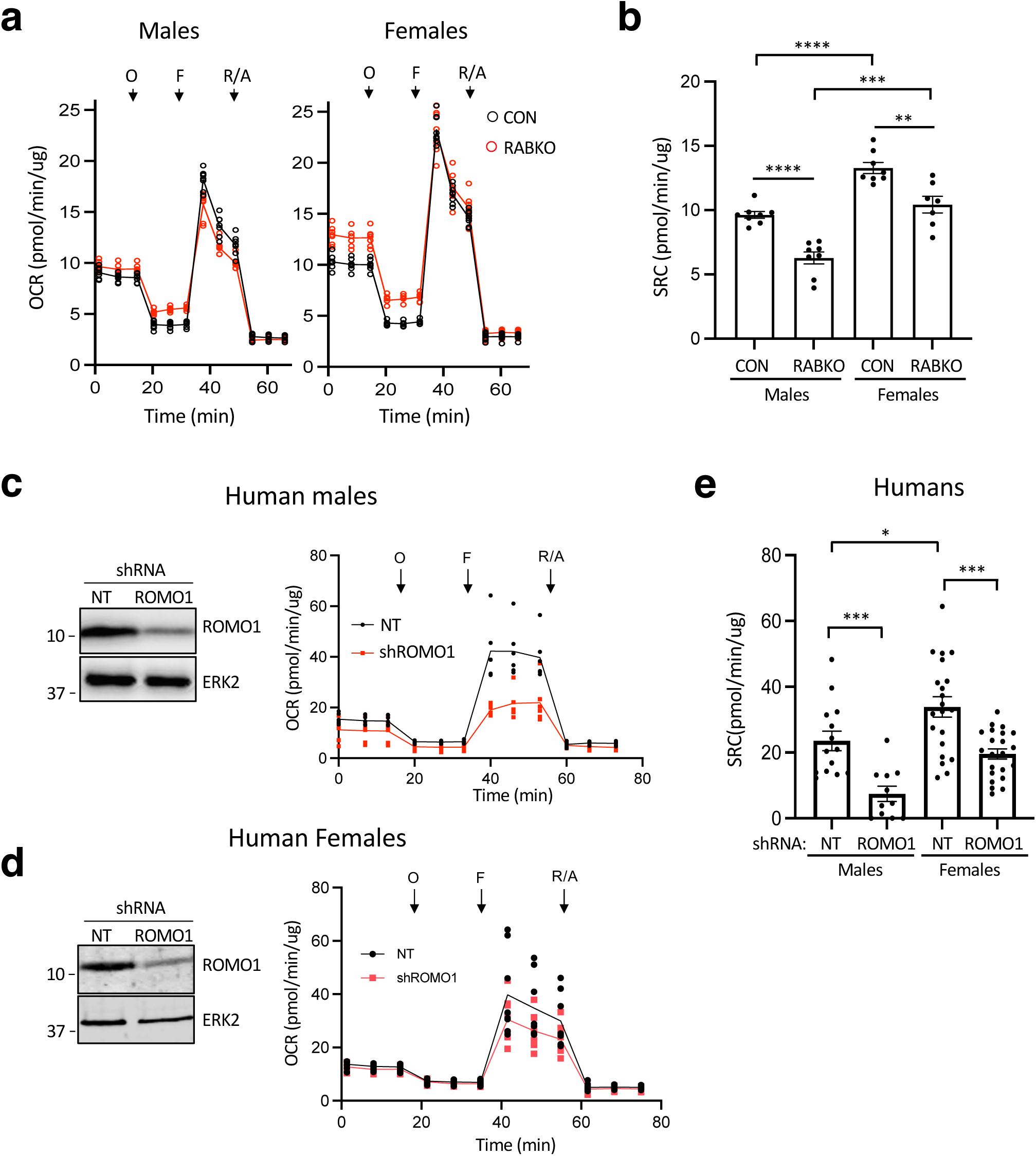
ROMO1 is required for SRC in mouse islets. a. Seahorse assay showing OCR in dissociated islet cells isolated from male and female CON (black lines) and RABKO mice (red lines). Treatment with oligomycin (O), FCCP (F), rotenone/antimycin A (R/A) are indicated. b. Barplot of SRC from Seahorse assay data shown in panel a. P-values ** < 0.01, *** < 0.001, **** < 0.0001. c. Representative seahorse assay showing oxygen consumption rate and loss of SRC in male human islet cells following ROMO1 silencing. Treatment with oligomycin (O), FCCP (F), and rotenone/antimycin A (R/A) are indicated. d. Representative seahorse assay showing oxygen consumption rate and loss of SRC in female human islet cells following ROMO1 silencing. Treatment with oligomycin (O), FCCP (F), and rotenone/Antimycin A (R/A) are indicated. e. Barplot of SRC from Seahorse assays of male (n=3) and female (n=3) human islets following ROMO1 silencing. P-values: ***Male non-targeting shRNA (NT) vs. shROMO1, P < 0.001; ***Female non-targeting shRNA (NT) vs. shROMO1, P < 0.001; *Male CON vs. Female CON, P < 0.05.

### Aging drives glucose intolerance in RABKO mice

The transition from prediabetes to T2D is characterized by a loss of beta cell function [31–33]. Aging is associated with increased incidence of chronic inflammation and insulin resistance, known risk factors for T2D [34]]. To evaluate the potential contributions of aging to the RABKO phenotype, we initiated ROMO1 deletion in male and female mice at 8 weeks of age then analyzed glucose tolerance and islet function at or beyond 1 year of age. Islet morphology and insulin staining in aged mice were indistinguishable from younger mice and between sexes (Supplementary Figure 4). Glucose tolerance tests revealed pronounced glucose intolerance and reductions in plasma insulin in RABKO males at 52 weeks of age compared to controls (Figure 6a,b), such that a 50% lower glucose bolus was required to monitor glucose excursions. Female RABKO mice were also glucose intolerant with reductions in plasma insulin at 15- and 30 minutes post injection of glucose, indicating that female RABKO mice lose the capacity to compensate for loss of ROMO1 as they age. While young and old male RABKO mice showed a complete loss of SRC compared to control male mice (Figure 6e), only young female RABKO mice maintained SRC at the levels of control mice, with female RABKO mice at 52 weeks showing loss of SRC (Figure 6f). The loss of glucose tolerance and SRC in aged RABKO females was coincident with a reduction in mtDNA (Figure 6g).

**Figure 6.**
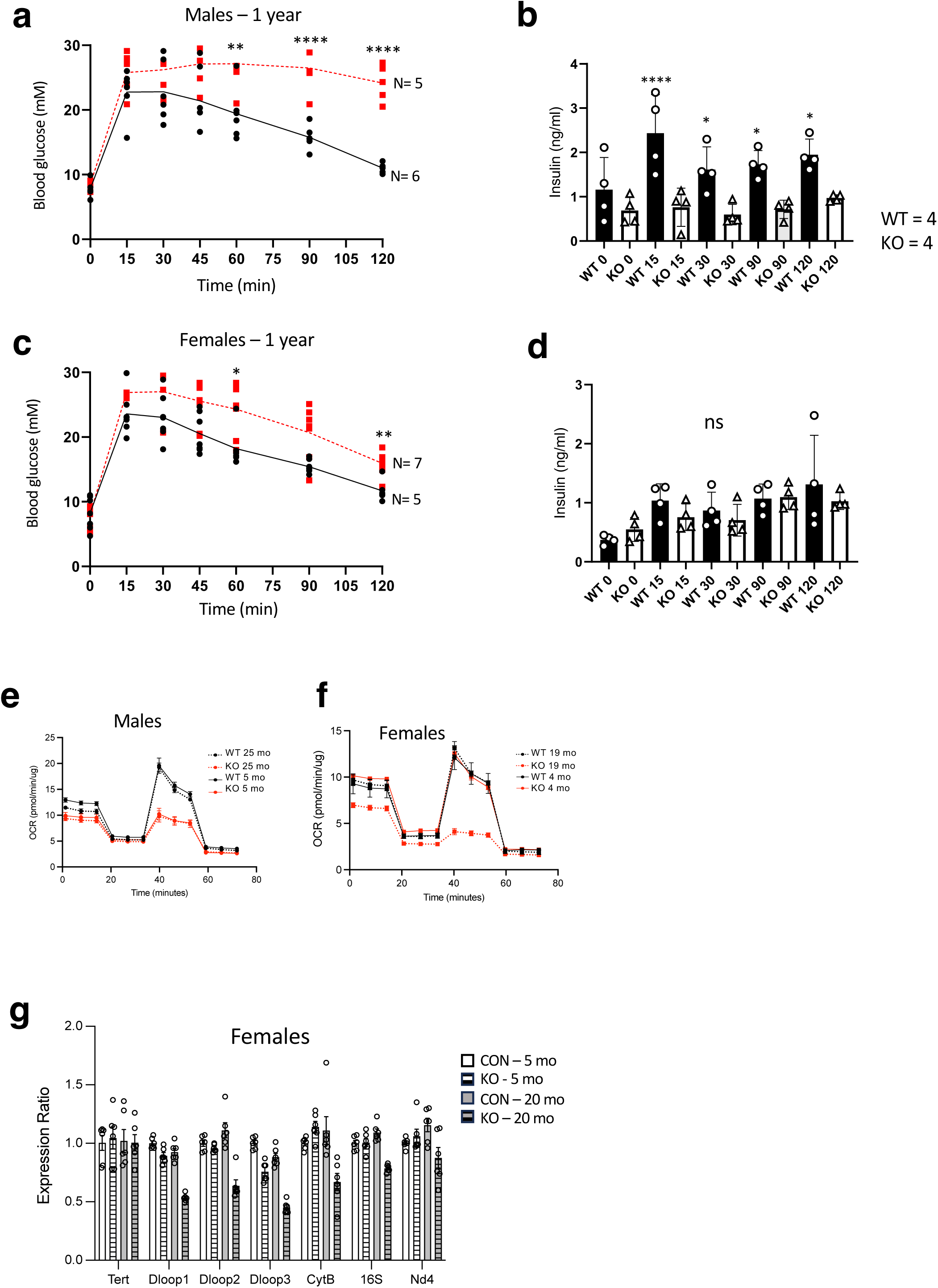
Aging results in loss of SRC and glucose intolerance in females. a. IPGTT assay in CON and RABKO 52-week-old male mice after injection of 1g glucose/kg body mass. P-values = **60 min = 0.007, ***90 min = 0.0008, ***120 min = 0.0005. b. Plasma insulin levels in CON (black bars) and RABKO (white bars) 52-week-old male mice during IPGTT glucose challenge from a. P-values = **15 min = 0.00003, **30 min = 0.0168, *90 min = 0.0168, *120 min = 0.0168. c. IPGTT assay in CON and RABKO 52-week-old female mice. P-values = *15 min 0.025, *30 min 0.034, *45 min 0.025, **60 min 0.007, ***90 min 0.0008, ***120 min 0.0005. d. Plasma insulin levels in CON (black bars) and RABKO (white bars) 52-week-old female mice during IPGTT glucose challenge from c. All data N.S. e. Seahorse assay showing OCR (left) and SRC (right) in dissociated islet cells isolated from 5 month (solid lines) or 25-month-old (dotted lines) male CON (black lines) and RABKO mice (red lines). Treatment with oligomycin (O), FCCP (F), rotenone/antimycin A (R/A) are indicated. f. Seahorse assay showing OCR (left) and SRC (right) in dissociated islet cells isolated from 4-month-old (solid lines) and 19-month-old (dotted lines) female CON (black lines) and RABKO mice (red lines). Treatment with oligomycin (O), FCCP (F), rotenone/antimycin A (R/A) are indicated. g. qPCR data of mitochondrial DNA in female CON and RABKO mice at 5 and 20 months of age. Relative levels of mtDNA genes Dloop1, Dloop2, Dloop3, CytB, mito16S, and Nd4 in islets from CON and RABKO female mice were normalized to level of the nuclear DNA gene, Tert.

### ROMO1 and Complex II are essential for SRC

Under normal conditions, most electrons enter the electron transport chain at complex I, contributed by NADH, with the remaining entering an auxiliary path at complex II, known also as succinate oxidoreductase or succinate dehydrogenase (SDH) of the TCA cycle [35]. Recent work has implicated mitochondrial complex II as a source of SRC, as under stress, mitochondria increase their respiratory output by increasing electron flow through this auxiliary path [36]. Consistent with this, complex II is critical both in development and in aging, where loss of complex II respiration results in succinate accumulation and ROS generation, and drives age-related diseases, including diabetes [37]. Furthermore, loss of SRC has been linked to respiratory activity at complex II [38]. To determine the effect of loss of ROMO1 on mitochondrial respiratory complexes, we silenced ROMO1 in U2OS cells and monitored oxygen consumption in the presence of fuels specific to complexes I, II, III, and IV by Seahorse assay (Figure 7a). While cells lacking ROMO1 displayed normal oxygen consumption in the presence of pyruvate (complex I), G3P (complex III), and TMPD+ascorbate (complex IV), respiration at complex II was uniquely defective. We then compared OCR in the presence of succinate in male and female control and RABKO islets and observed a specific reduction in respiration in RABKO islets from male, but not female mice (Figure 7b). Taken together, these data indicate that complex II defect in the absence of ROMO1 underlies the loss of glucose coupling and susceptibility to diabetes in RABKO mice.

**Figure 7.**
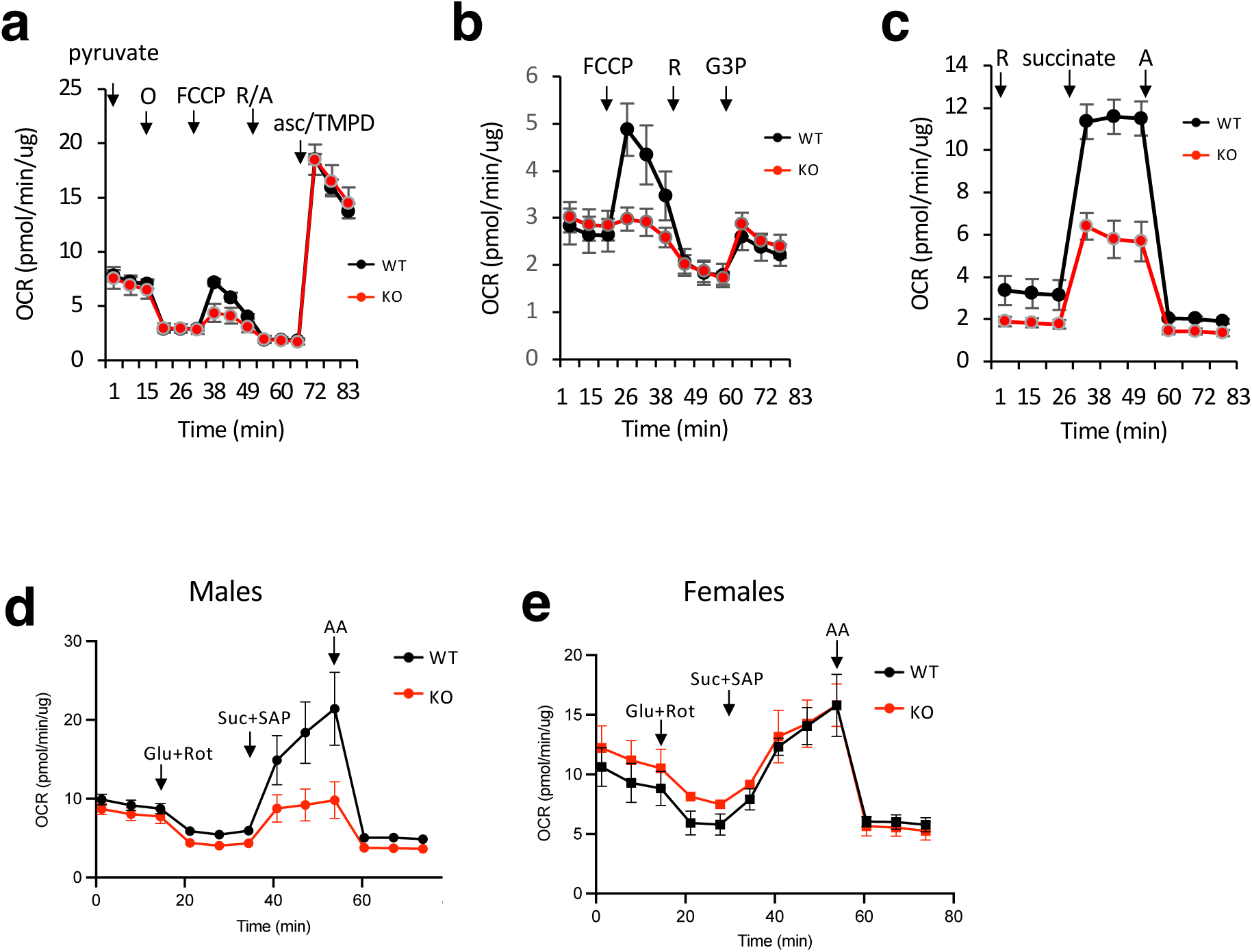
ROMO1 is required for complex II activity. a. Seahorse assay showing OCR in control cells (black lines) and cells after ROMO1 silencing (red lines) in the presence of pyruvate. Treatment with oligomycin and FCCP is shown. Addition of rotenone and antimycin A to inhibit complexes I and III prior to addition of complex IV substrate ascorbate/TMPD is shown. c. Seahorse assay showing OCR resulting from activity at complex III (glyceraldehyde 3-phosphate, G3P) in control (black line) and after ROMO1 silencing (red line). c. Seahorse assay showing OCR resulting from activity at complex II (succinate) in the presence of complex I inhibitor rotenone in control (black line) and after ROMO1 silencing (red line). d. Seahorse assay showing OCR in 16-week-old male control mice (black line) and RABKO mice (red line) resulting from activity at complex II after addition of succinate (Suc) in cells permeabilized with saponin (SAP). Treatment with glucose and rotenone (Glu+Rot) and antimycin shown. Seahorse assay showing OCR in 16-week-old female control (black line) and RABKO (red line) mice resulting from activity at complex II after addition of succinate in the presence of complex I inhibitor rotenone.

## Discussion

Mitochondrial SRC is essential for the generation of ATP by oxidative phosphorylation under increased demand [36]. In this study, we used a null allele of *Romo1* to evaluate the effect of defective mitochondrial SRC *in vivo* and show that ROMO1 is essential for early embryonic development in mice.

It is noteworthy that *Romo1* null mice die peri-implantation (E4-E8), earlier than that of mice lacking *Mfn1*, *Mfn2*, or *Opa1*, which like *Romo1*, are necessary for mitochondrial fusion [18, 39–41]. ATP levels in control and ROMO1 silenced U2OS cells were unchanged, indicating loss of ROMO1 is not required for maintenance of steady-state cellular energy levels in this context (Supplementary Figure 1). The timing of the embryonic lethality also precedes that of genes involved in fragmentation (Drp1 at E11.5[42]), oxidative phosphorylation (Cytc, E10.5 [43]) or mtDNA maintenance (Rnash1 [44], Mterf3[45], Tfam [46], Polg [47]). This suggests that, in addition to the regulation of mitochondrial fusion [18], ROMO1 may serve a more primordial bioenergetic role essential for embryonic development in mice.

How ROMO1 functions has been controversial. Richter and colleagues have reported a global loss of complex IV in ROMO1 null cells [16], however, complex IV-dependent respiration was normal in RABKO islets. It has been reported that ROMO1 governs the selective import of mitochondrial protein targets, including the YME1L protease [16], an important regulator of mitochondrial quality control. We do not observe a global reduction in mitochondrial respiration or a complex IV-specific defect in ROMO1 KO cells and tissues, nor do we see a defect in import of YME1L (not shown). We ascribe these differences to the systems/approaches used (HEK293T cell line *vs*. primary mouse tissues), and to different gene ablation strategies. Our gene targeting approach was designed to avoid exon 1 of Romo1 as it overlaps with the *Nfs1* gene that is transcribed from the opposite strand, and we verified that *Nfs1* expression was normal in mouse tissue following Cre-mediated *Romo1* deletion. *Nfs1* encodes a cysteine desulfurase, that is required for the assembly of mitochondrial iron-sulfur clusters, ancient protein modules that serve essential roles in enzyme catalysis and electron transport, including ETC complexes I and III [48]. Consistent with the reported growth defect in CRISPR-Cas9 *Romo1* deletion cells that was only partially relieved by the reintroduction of ROMO1 [16], it is conceivable that the CRISPR-Cas9 ROMO1 KO cells may be a compound knockout of *Romo1* and *Nfs1*.

Sex differences in homeostatic physiology are being recognized as the basis for differences in both pathology and treatment responses in specific brain regions and in stress-based pathophysiology, and this is true for insulin secretion in both rodent models and humans [49, 50]. Our results revealed a sexual dimorphism in young mice with respect to ROMO1 in the β cell being essential for the maintenance of euglycemia only in males. At four months of age, male RABKO mice show glucose intolerance and insulin secretion defects, while female RABKO mice secrete insulin normally and display normal fasting glucose and glucose tolerance. We have reported that ablation of ROMO1 leads to loss of SRC in cultured cells and in the mouse heart [18, 51], and our results in the mouse beta cell indicate that ROMO1 is required for SRC in diverse settings. Interestingly, our analysis of SRC in islets revealed that deletion of ROMO1 lowered SRC in both male and female mice, islets from male mice and humans have a significantly lower SRC setpoint than females, indicating that higher SRC is a general property of female beta cells/islets and that females have both ROMO1-dependent and -independent mechanisms for maintaining SRC.

The morphology of mitochondria and mtDNA maintenance in RABKO mice is also sexually dimorphic. Mitochondria from males lacking ROMO1 are swollen and show disorganized and even floating cristae, which may be driven by the greater loss of mtDNA we observed in males. How might the loss of ROMO1 lead to mitochondrial morphology changes? Membrane curvature associated with cristae junctions is thought to be established by dimerization of ATP synthase (complex V) and of the MICOS complex, which establishes a boundary between the IMM that are dedicated to cristae vs. non-cristae regions [52, 53]. ROMO1 co-precipitates with mitofilin [18], the core component of the MICOS complex, and with OPA1, the GTPase required for IMM fusion. In the absence of ROMO1, OPA1 processing is abnormal [18] and likely contributes to dysfunctional IMM dynamics. However, our data demonstrating loss of SRC in the absence of ROMO1 prompt a model whereby loss of bioenergetic capacity lies upstream of dysfunction in MICOS and OPA1. That young female mice show a reduction in SRC yet maintain normal cristae morphology and mtDNA levels indicate that the bioenergetic defect at complex II occurs prior to any ultrastructural changes. However, the Yoo group reported that ROMO1 can function as a nonselective cation channel *in vitro* [17], and proposed that six ROMO1 molecules could form a hexameric pore-forming unit, a structure that could promote the membrane curvature required to generate or maintain cristae folds. Future work will a potential role of ROMO1 in regulating ion transport in the beta cell.

We found that deletion of ROMO1 in cultured cells and primary islets ablates mitochondrial respiration, specifically at complex II/SDH. Complex II displays several characteristics that make it unique from complexes I, III, IV and V. First, the succinate oxidoreductase activity of complex II comprises a step in the TCA cycle, thus coupling oxidation to electron transport in diverse cell types [54]. Second, all four of its subunits (SDHA, SDHB, SDHC, and SDHD) are nuclear-encoded and thus they are not subject to ROS-mediated mtDNA damage. Third, complex II is clustered near cristae junctions and does not line the entire length of cristae [55, 56. Our data indicate that reducing equivalents generated at complex II, and not through the canonical complex I-III-IV pathway, is essential for SRC and glucose-stimulated insulin secretion, for which there is precedence {Edalat, 2015 #1328].

Aging is a risk factor for glucose intolerance in both male and female RABKO mice, as glucose tolerance and insulin secretion are negligible in males and significantly impaired in females. Interestingly, female mice > 1 year of age develop glucose intolerance that is coincident with complete loss of SRC and reductions in mtDNA, indicating that aging in females is sufficient to unmask a requirement for ROMO1 in the beta cell. Significant reductions in insulin secretion are observed with aging in humans [57], an effect that may be attributed to sex-specific genomic methylation patterns [58]. As loss of SRC is a hallmark of aging cells [20], the lower SRC setpoint in male islets may, in part, explain the preferential loss of insulin secretion in human male populations [58, 59], and suggest that beta cells in males may be more susceptible to genetic or environmental triggers that impair SRC. Our data indicate that strategies to promote ROMO1 activity and further exploration of the molecular basis for the higher SRC in females would help to enhance glucose coupling, insulin secretion and glucose tolerance in settings where insulin levels are low.

## Abbreviations

IMM: inner mitochondrial membrane
OMM: outer mitochondrial membrane
OPA1: Optic atrophy type 1
RABKO: ROMO1 adult beta cell knockout
ROMO1: Reactive oxygen species modulator 1
ROS: reactive oxygen species
SDH: succinate dehydrogenase
SRC: spare respiratory capacity
TCA: tricarboxylic acid
T2D: Type 2 diabetes
YME1L: YME-like 1 ATPase

## Acknowledgements

LW, CI, QY, ZC, and ACHN performed experiments and edited manuscript. CR, MP, and SPY performed experiments. RAS designed the project and wrote the manuscript. We would like to acknowledge Dalia Barayan and Carly Knuth for helpful comments, Lindsey Fiddes at the University of Toronto Microscopy Imaging Lab and Ali Darbandi at Hospital for Sick Kids, Toronto, for EM sample processing and imaging, and Judy Cathcart University Health Network, Division of Biophysics and Bioimaging for imaging of stained pancreatic sections. The authors declare no conflict of interest. The project was supported by Canadian Institutes of Health Research Project Grant 142352 and Diabetes Canada End Diabetes:100 Award OG-3-21-5584-RS to RAS. RAS is the guarantor of this work and, as such, had full access to all the data in the study and takes responsibility for the integrity of the data and the accuracy of the data analysis. Some of the data in the manuscript has been presented at scientific meetings.

