## Supplemental Figures for "ROMO1 and Mitochondrial Complex II/SDH are Required for Spare Respiratory Capacity and Glucose Homeostasis in Mice"

### Supplementary Figures

**1**

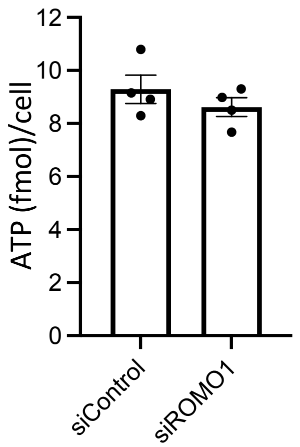


**2**

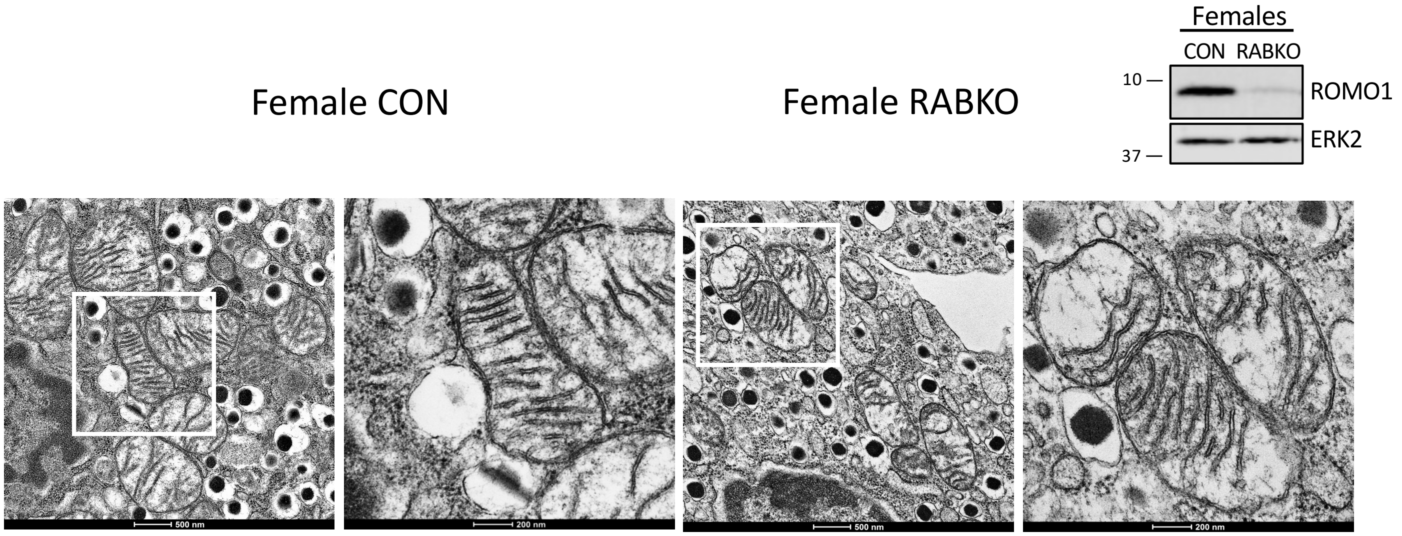


**Supplementary Figure Legends**

1. ATP levels in U2OS cells transfected with control non-targeting (CON) or ROMO1 RNAi duplexes were determined by mass spectrometry.
2. Left: Electron micrographs of beta cells from female *Romo1*$^{+/+}$*;Pdx1-CreERTam* (CON) and *Romo1*$^{fl/fl}$;*Pdx1-CreERTam* knockout (RABKO) mice at 12 week of age (4 weeks post-tamoxifen injection) showing normal cristae morphology in CON and RABKO female mice. Scale bar = 200 nm. Right: Western blot showing loss of Romo1 protein in islets from female RABKO mice.

### Supplementary Figures

**3**

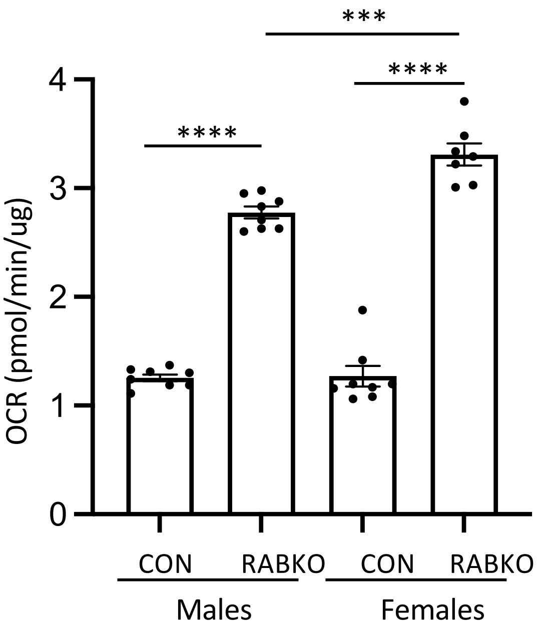


Proton Leak

**Supplementary Figure Legends**

3. Oxygen consumption attributed to proton leak in islets isolated from female and male WT and RABKO mice. P-values *** <0.001, **** < 0.0001

4

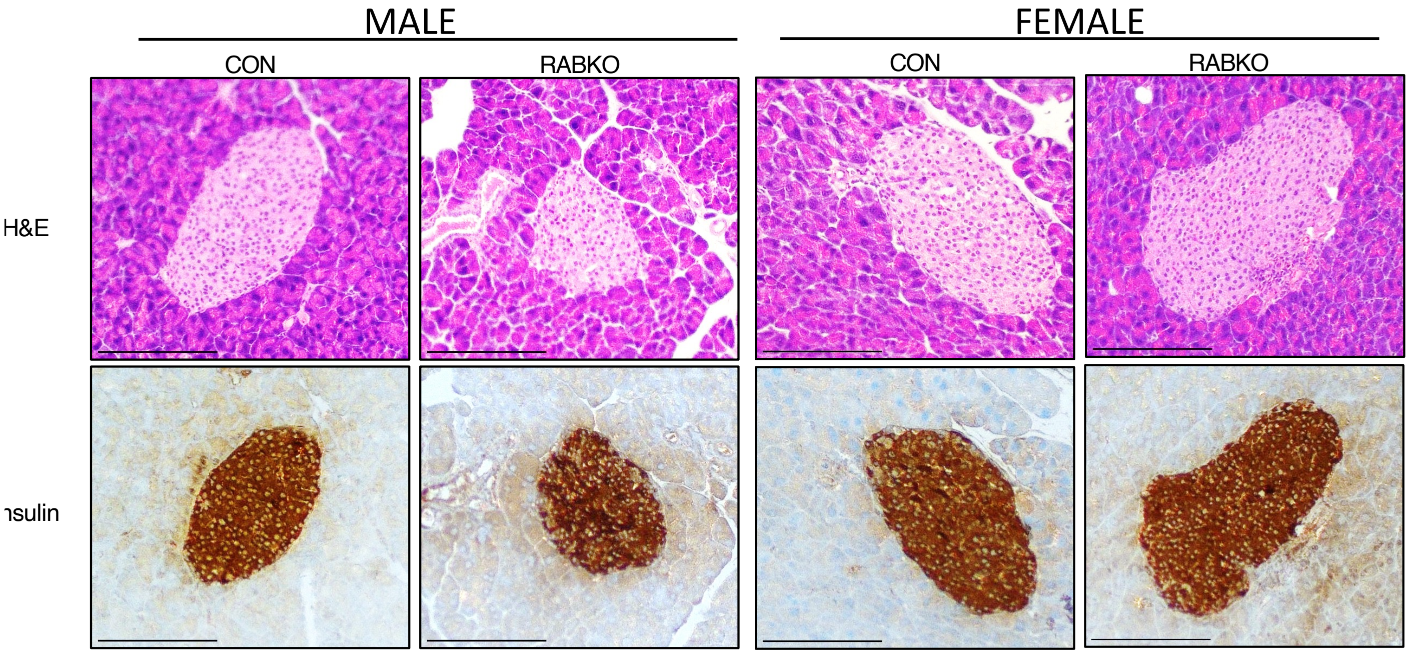


**Supplementary Figure Legends**

4. Hematoxylin and eosin micrographs (top) and insulin staining (bottom) in 1 year old male and female control and RABKO mice. Scale bars = 100 um.
